# Analyzing the heterogeneity of rule-based EHR phenotyping algorithms in CALIBER and the UK Biobank

**DOI:** 10.1101/685156

**Authors:** Spiros Denaxas, Helen Parkinson, Natalie Fitzpatrick, Cathie Sudlow, Harry Hemingway

## Abstract

Electronic Health Records (EHR) are data generated during routine interactions across healthcare settings and contain rich, longitudinal information on diagnoses, symptoms, medications, investigations and tests. A primary use-case for EHR is the creation of phenotyping algorithms used to identify disease status, onset and progression or extraction of information on risk factors or biomarkers. Phenotyping however is challenging since EHR are collected for different purposes, have variable data quality and often require significant harmonization. While considerable effort goes into the phenotyping process, no consistent methodology for representing algorithms exists in the UK. Creating a national repository of curated algorithms can potentially enable algorithm dissemination and reuse by the wider community. A critical first step is the creation of a robust minimum information standard for phenotyping algorithm components (metadata, implementation logic, validation evidence) which involves identifying and reviewing the complexity and heterogeneity of current UK EHR algorithms. In this study, we analyzed all available EHR phenotyping algorithms (n=70) from two large-scale contemporary EHR resources in the UK (CALIBER and UK Biobank). We documented EHR sources, controlled clinical terminologies, evidence of algorithm validation, representation and implementation logic patterns. Understanding the heterogeneity of UK EHR algorithms and identifying common implementation patterns will facilitate the design of a minimum information standard for representing and curating algorithms nationally and internationally.

## 1 Introduction

In the United Kingdom (UK), structured electronic health records (EHR) spanning primary care, hospital care, disease/procedure registries and death registries are used to create longitudinal disease phenotypes for observational research studies [Hemingway *et al*., 2018]. Through a process called *phenotyping*, researchers create algorithms which utilize multiple EHR sources to accurately extract information on diseases (e.g. status, onset and progression), lifestyle risk factors and biomarkers [Banda *et al*., 2018]. Phenotyping however is challenging due to the fact that EHR are fragmented, curated using different controlled clinical terminologies and collected for purposes other than research (e.g. reimbursement, audit) [Morley *et al*., 2014].

Phenotyping requires a significant amount of resources and mix of expertise, yet no common standard approach for defining, validating and ultimately sharing EHR phenotyping algorithms currently exists. In the UK, structured primary care EHR have been used in >1,800 peer-reviewed studies to date but only 5% of studies published sufficiently reproducible phenotypes [Springate *et al.,* 2014]. Defining a standardized format to represent EHR phenotypes will enable portability across data sources (and healthcare systems) and facilitate the systematic sharing of algorithms across the community [Mo *et al*. 2015].

Compared to the United States (US), the UK EHR research landscape differs in two important ways: 1) researchers can utilize multiple national EHR sources to create longitudinal ‘cradle to grave’ phenotypes [Kuan *et al.,* 2019], and 2) UK primary care EHR contain both healthy and unhealthy individuals which allow researchers to capture information on disease severity and progression over time. A recent systematic review identified 66 different definitions used to capture asthma status and exacerbations in research using UK EHR [Al Sallakh *et al*., 2017] demonstrating significant existing heterogeneity. While analyses have been undertaken in the US to characterize the heterogeneity of phenotyping algorithms [Conway *et al*., 2011], no such analysis has been carried out in the UK.

One of the aims of the newly-established national institute for health data science, Health Data Research UK (HDR UK, www.hdruk.ac.uk), is the creation of a national Phenomics Resource: an open-access online resource where EHR phenotypes can be deposited and curated. A critical first step in this process is to establish a minimum information standard for representing EHR phenotyping algorithms. This involves exploring and documenting the complexity, heterogeneity, design and implementation patterns of contemporary phenotyping algorithms in the UK. The concept of a minimum information standard has been used successfully in other biomedical disciplines, e.g. Minimum Information About a Microarray Experiment (MIAME) defines standards for reporting microarray experiments [Brazma *et al*., 2001]. Establishing a standardized method for representing phenotypes in the UK can potentially address these challenges and ensure compatibility with other international initiatives such as eMERGE and PCORNet [Fleurence *et al*. 2014; Gottesman *et al*. 2013].

## 2. Aims

Despite the widespread use of UK EHR data sources for research, contemporary research resources utilize different approaches for algorithm creation, curation and validation. The aims of this study were to: a) identify and characterize the structural components, implementation logic and heterogeneity of rule-based algorithms defining diseases, lifestyle risk factors and biomarkers in structured national EHR in the UK utilized by contemporary research resources, and b) propose a minimum information standard to represent UK EHR phenotyping algorithms.

## 3. Methods

We identified, downloaded and reviewed published phenotyping algorithms for diseases, biomarkers and lifestyle risk factors from two large-scale contemporary UK research resources: UK Biobank^1^ and CALIBER^2^.

The UK Biobank [Sudlow *et al*., 2015] is a prospective cohort study of 500,000 (aged 40-69 at recruitment) adults recruited in England, Scotland and Wales from 2006-2010. For each participant, deep phenotypic and genotypic information is available including biomarkers in blood and urine, imaging (brain, heart, abdomen, bone, carotid artery), lifestyle indicators, pathophysiological measurements and genome-wide genotype data. Follow-up for health outcomes is enabled by hospital EHR (Hospital Episode Statistics (HES) in England, Patient Episode Data Warehouse in Wales and Scottish Morbidity Registry in Scotland) and linkages to primary care EHR are underway. CALIBER [Denaxas *et al*., 2012; Denaxas *et al*., 2019] is a research resource consisting of algorithms, tools and methods for structured EHR linked across primary care (Clinical Practice Research Datalink, CPRD), hospital care (HES) and a mortality data (Office for National Statistics, ONS) in the UK.

In the UK, national EHR are recorded using controlled clinical terminologies where terms are assigned at variable timepoints i.e. in UK primary care the physician records terms in real time during the consultation with the patient whereas in hospital care terms are retrospectively entered into databases by trained coders and data selected for billing purposes. We identified and counted the number of ontology terms each algorithm utilizes from five controlled clinical terminologies which are widely used in the UK: a) Read (primary care, subset of SNOMED-CT), b) International Classification of Diseases 9th and 10th Revision (ICD-9, ICD-10, secondary care diagnoses and cause of mortality), c) OPCS Classification of Interventions and Procedures (OPCS-4, hospital surgical procedures, analogous to the Current Procedural Terminology ontology used in the United States), and d) the Dictionary of Medicines and Devices (DM+D) which is used to record primary care prescriptions. Terms were automatically extracted from documents and counted using regular expressions in Python 3.6^3^. We manually extracted and counted terms across five randomly chosen algorithms to verify the automatically-generated counts.

EHR phenotype validation is a critical process guiding the subsequent use of algorithms and we were interested in what types, if any, of evidence were available to external researchers. We classified the available material into six non-overlapping categories which encapsulate all potential approaches for obtaining validity evidence (adapted from [Denaxas *et al,* 2019] and recorded as used/not used):

- **Aetiological**: Are the prospective associations with risk factors consistent with previous published evidence from both EHR and non-EHR studies?
- **Prognostic**: Are the risks of subsequent events plausible and consistent with existing domain knowledge?
- **Case-note review**: What is the positive predictive value (PPV) and the negative predictive value (NPV) when comparing the algorithm with clinician-led review of case notes, self-reported information or a suitable “gold standard” source?
- **Cross-EHR-source concordance**: To what extent is the phenotype concordant across EHR sources?
- **Genetic**: Are the observed genetic associations plausible and consistent in terms of magnitude and direction of association with associations reported from non-EHR studies?
- **External populations**: Has the algorithm been evaluated in different countries or external sources?

For each algorithm, we documented the EHR sources the phenotype is derived from (i.e. primary care, hospital care, mortality register). We extracted information on the representation components of phenotypes e.g. the presence of tabular data and the use of a flowchart (or other graphical presentation). We extracted and categorized information on the different types of implementation logic, temporality and algorithm implementation patterns (Table 1), partially based on previous research in the US [Conway *et al.,* 2011].

**Table 1:** Characteristics of implementation logic, temporality and algorithmic implementation features extracted and analyzed from phenotyping algorithms in the UK Biobank and the CALIBER resources. AF Atrial Fibrillation; BMI Body Mass Index; BP Blood Pressure; DBP Diastolic Blood Pressure; DVT Deep Vein Thrombosis; PVD Peripheral Vascular Disease; PE Pulmonary Embolism; HT Hypertension; HF Heart Failure; mmHg millimeter of mercury; SBP Systolic Blood Pressure.

| Concept | Definition | Example |
| --- | --- | --- |
| **Simple Boolean** | Simple Boolean statements e.g. “AND”, “OR” | PVD diagnosis during a primary care consultation OR diagnosis of leg or aortic embolism or thrombosis during a hospitalization |
| **Complex Boolean** | Nested statements with multiple layers? | IF patient = diabetic: HT threshold: SBP ≥140 mmHg OR DBP ≥90 mmHg ELSE: threshold SBP ≥150 mmHg OR DBP ≥90 mmHg |
| **Negation** | Are negation statements used? | No AF diagnosis term is present, but the patient record includes a warfarin prescription in the absence of prior DVT or PE, or a digoxin prescription but no HF |
| **Temporal (simple)** | Temporal proximity future or past | Iron deficiency anaemia record in primary care OR hospital AND endoscopy in 30 days |
| **Temporal (complex)** | Complex temporal rules. multiple logic layers | ≥3 high SBP/DBP readings within 1-year OR ≥2 high SBP/DBP readings in a 6-month period |
| **Biomarker** | Evidence from continuous measurement | Presence of a positive rheumatoid factor test or anti-cyclic citrullinated peptide antibody test after a rheumatoid arthritis diagnosis |
| **Complex calculation** | Calculation e.g. unit conversion | Calculate average BMI in consultation, exclude measurements $<10 \text{ kg/m}^2$ or $>80 \text{ kg/m}^2$ |

## 4. Results

We identified and reviewed 70 EHR phenotyping (Table 2) algorithms available from the UK Biobank (n=19) and the CALIBER resource (n=51). The majority of phenotyping algorithms were created to ascertain disease status (n=54) (e.g. heart failure [Gho *et al*. 2018; Uijl *et al*. 2019], depression [Daskalopoulou *et al*. 2016]), ten algorithms were created to extract information on biomarkers (e.g. heart rate [Archangelidi *et al*. 2018], blood pressure [Rapsomaniki *et al*. 2014]) and six algorithms were used to identify lifestyle risk factors (e.g. alcohol [Bell *et al*. 2017], smoking [Pujades-Rodriguez *et al*. 2015]).

**Table 2.**
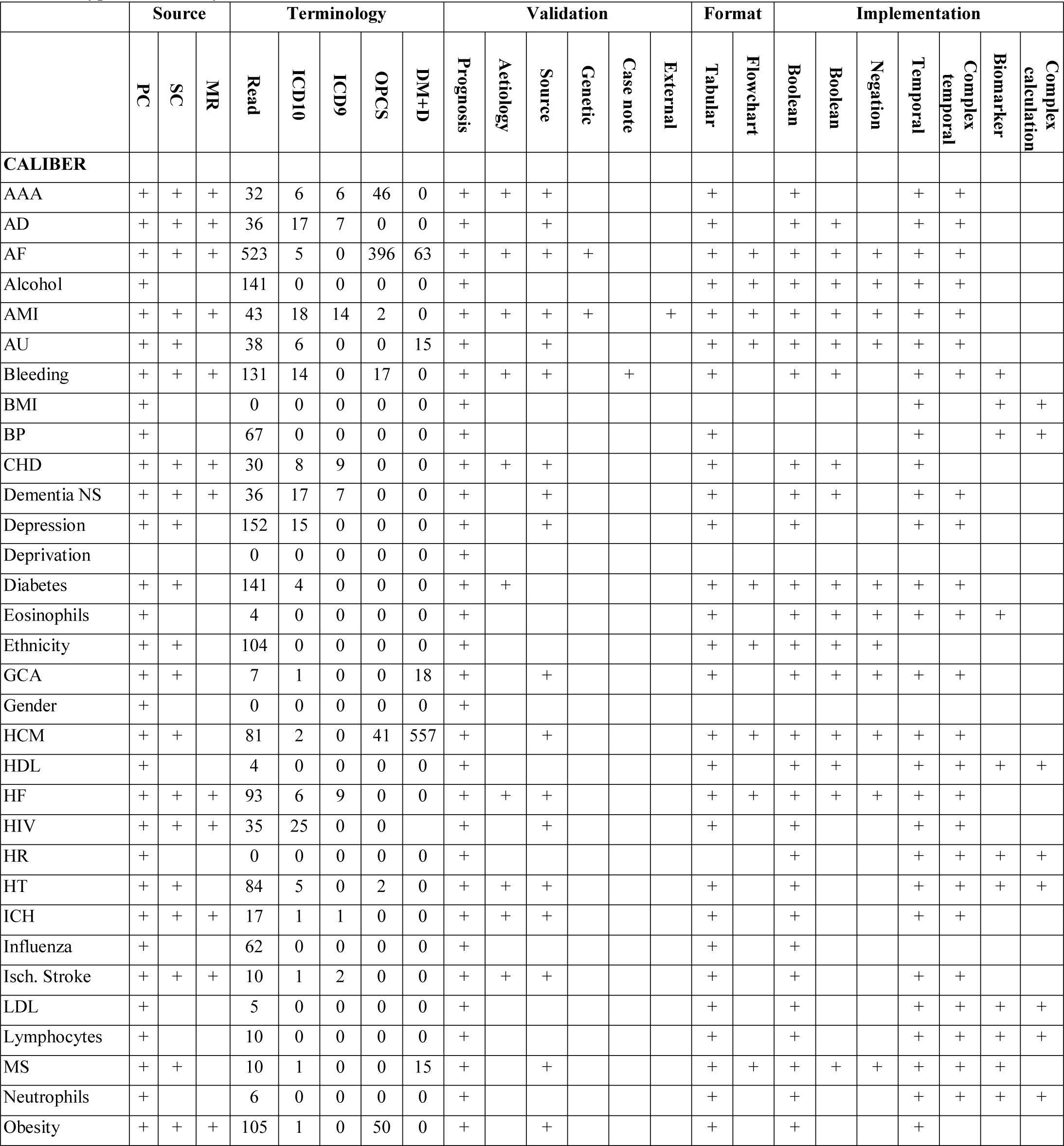
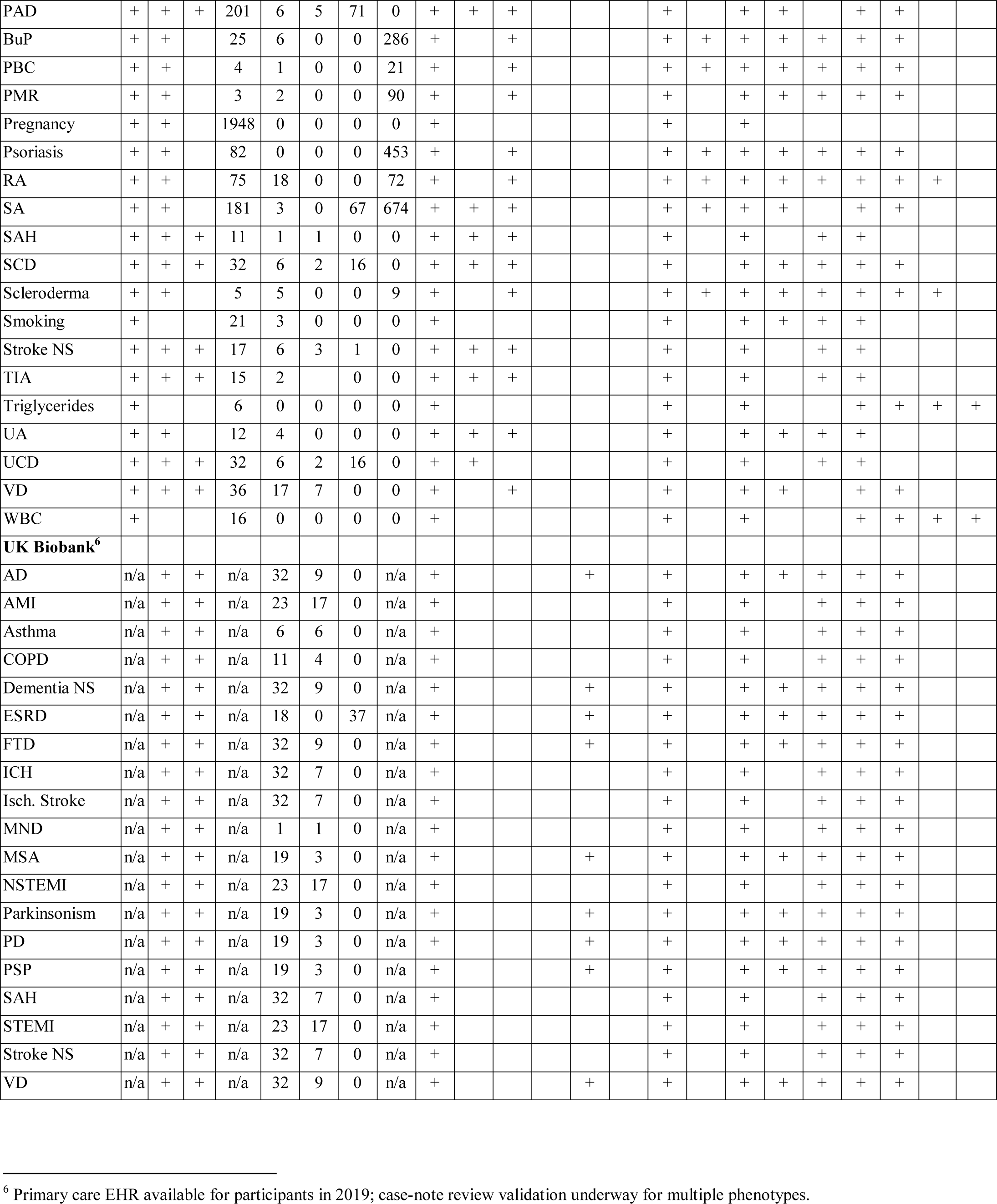
Information on EHR data sources, controlled clinical terminologies, available evidence of algorithm validation, algorithm representation format and implementation logic patterns from UK Biobank and CALIBER EHR phenotype algorithms. AAA Abdominal Aortic Aneurysm; AD Alzheimer’s Disease; AF Atrial Fibrillation; AMI Acute Myocardial Infarction; AU Autoimmune Uveitis; BMI Body Mass Index; BP Blood Pressure; BuP Bullous Pemphigoid; CHD Coronary Heart Disease; FTD Frontotemporal dementia; GCA Giant Cell Arteritis; HCM Hypertrophic Cardiomyopathy; HDL High Density Lipoprotein cholesterol; HF Heart Failure; HIV Human Immunodeficiency Virus; HR Heart Rate; HT Hypertension; ICH Intracerebral Haemorrhage; LDL Low Density Lipoprotein cholesterol; MS Multiple Sclerosis; NS Not Specified; PAD Peripheral Arterial Disease; PBC Primary Biliary Cirrhosis; PMR Polymyalgia Rheumatica; RA Rheumatoid Arthritis; SA Stable Angina; SAH Subarachnoid Haemorrhage; SCD Sudden Cardiac Death; TIA Transient Ischaemic Attack; UA Unstable Angina; UCD Unheralded Coronary Death; VD Vascular Dementia; WBC White Blood Cell Count; COPD Chronic Obstructive Pulmonary Disease; ESRD End Stage Renal Disease; MND Motor Neuron Disease; PD Parkinson’s Disease and Parkinsonism; MSA Multiple System Atrophy; PSP Progressive Supranuclear Palsy; STEMI ST-Elevation AMI; NSTEMI Non-ST Elevation AMI

All but one CALIBER phenotyping algorithm (n=50) used information from primary care EHR with the exception of socioeconomic status which was defined using the Index of Multiple Deprivation (IMD) provided by the ONS. Algorithms defining biomarker measurements (e.g. white blood cells, heart rate) were based on primary care EHR entirely while approximately half of the algorithms ascertaining disease status (n=19 of 35) combined information across all three EHR sources. All currently available UK Biobank algorithms (n=19) combined information recorded during the baseline assessment (data not shown), diagnoses and/or surgical procedures recorded during hospitalization and information based on the underlying (or secondary) cause of death which is recorded in the national mortality register. Primary care linkages in UK Biobank are still underway and as a result none of the currently available algorithms utilized information from primary care EHR. However, primary care information for just under half of the cohort (n=230,000) will be made available for UK Biobank researchers in June 2019. Algorithms incorporating primary care data for the conditions already covered have been or are being developed [Wilkinson *et al* 2019]. Along with a range of additional algorithms expanding the range of health outcomes available, they will be available from UK Biobank later in 2019. Overall, based on current publicly available information from CALIBER and UK Biobank, 75% (n=66) of algorithms used data from secondary care EHR and 45% (n=49) used information available in the death registry.

The most widely-used clinical terminology was Read with 4,729 (non-unique) terms used across all algorithms while the second highest number of terms was derived from the DM+D with 2,273 (non-unique) terms used to record prescriptions in primary care EHR. Four algorithms (body mass index, socioeconomic deprivation, sex, heart rate) did not use any terms across any terminology systems and were based on information which is derived from a structured field of the EHR or externally linked such as in the case of IMD. The atrial fibrillation algorithm used the highest number of clinical terms (n=987) while across all algorithms the pregnancy phenotype used the highest number of Read codes (n=1,948). ICD-9 was the terminology least used: in the UK Biobank it is used for recording diagnoses in older Scottish hospital records and in CALIBER it is used to record the cause of death prior to 1997. Algorithms defining biomarkers contained the lowest number of terminology terms as they relied on structured data fields combined with a small number of diagnosis terms to denote the type of test (e.g. Read code “42K..00 Eosinophil count”).

With regards to algorithm implementation logic, 66 (93%) of algorithms used Boolean statements, usually to identify the presence of one or more diagnosis codes in a patient’s EHR. Where Boolean statements were deployed, in nearly half of the cases these were complex and involved either a series of nested statements or joined information across multiple sources, for example in the UK Biobank where information is derived from self-reported, hospital and mortality sources and events are further stratified as ‘prevalent’ (first reported prior to recruitment) or ‘incident’ (first reported after recruitment). A similar pattern of logic was observed with regards to temporality where 66 algorithms utilized temporal rules and almost always this included more complex statements and restrictions. Finally, approximately half (n=43) of the algorithms used negation. Only ten algorithms (16%) included more complex calculations, usually to calculate the mean of multiple measurements on the same day or to harmonize units for laboratory measurements to a common format.

Prognostic 86% (n=66) and cross-source concordance 54% (n=43) validation approaches where the most widely-used algorithm evaluation approaches. The least-widely used validation approach was expert case note review, although this type of validation has been completed for a few UK Biobank algorithms, including dementia and its subtypes [Wilkinson *et al*, 2019], and is underway for several others. Most (93% [n=66]) of the algorithms used data stored in tabular format since tables are predominantly used to store lists of controlled clinical terminology terms. Only 25% (n=15) of algorithms included a graphical representation of the algorithm using a flowchart and all algorithms included a textual description of the algorithm components.

## 5. Discussion

In this study we downloaded and reviewed 70 EHR phenotyping algorithms from two large-scale, national research resources in the UK. We reviewed algorithms in terms of EHR data sources, controlled clinical terminologies used, available evidence of algorithm validation, algorithm representation formats and implementation logic patterns.

Similar to findings from US studies, we discovered that UK EHR algorithms make extensive use of Boolean statements and temporal logic. When these are used, they are often complex i.e. combining multiple nested Boolean layers of logic and defining temporal proximity rules within them. This is expected given that algorithms utilize multiple sources of information and include evidence from primary care and hospital care (or self-reported information in the case of the UK Biobank). Algorithms defining disease status were the most frequent and complex algorithms reviewed and utilized the greatest number of terms from controlled clinical terminologies. Negation was another major component of algorithms and is often used to exclude concomitant diagnoses or procedures when trying to ascertain diseases based on secondary information (e.g. ascertaining AF cases based on a prescription of digoxin but excluding patients which are diagnosed with HF).

The Read clinical terminology was the most popular terminology used with the highest number of terms per phenotype. These findings are expected as Read contains a significant amount of duplication internally due to synonym terms which can be potentially utilized. Additionally, the clinical concepts contained within Read subsume the concepts across all other terminologies i.e. Read contains terms for diagnoses, symptoms, laboratory tests, prescriptions and procedures. UK primary care clinical coding is currently transitioning to SNOMED-CT which should provide a more streamlined set of terms to be used.

In terms of validation, we observed a significant level of heterogeneity with approaches seeking to evaluate and replicate previously reported aetiological and prognostic estimates from non-EHR studies being the most popular. The presentation of the evidence however does not follow a common standard and sometimes only included references to published research rather than a more structured abstract of the main findings of the analyses. In contrast with the US, expert review of case records was the least frequently used approach for evaluation due to the fact that large scale corpuses of medical text do not exist in the UK owing to information governance restrictions and the technical challenges of integrating such data since they are held in a wide range of formats by multiple different NHS organisations. For similar reasons, none of the algorithms reviewed utilize medical text and natural language processing approaches to extract information from medical notes which is prevalent in some clinical specialties such as mental health [Wu *et al*. 2018].

Significant heterogeneity was also observed in terms of representation. UK Biobank algorithms were curated in individual PDF files^4^ and included extended information on the goal of the algorithm and useful background knowledge and references. In contrast, CALIBER phenotypes were stored in an online, openly-available Portal^5^, spanned multiple pages and did not include much background information. Flowcharts or similar graphical representations were not widely-used and while they are not machine-readable, they can potentially minimize errors during translation of the algorithm to machine code.

Our study has potential limitations. We reviewed algorithms from only two UK sources. While other UK initiatives exist, they tend to focus on curating lists of controlled clinical terminology terms (referred to as *codelists*) rather than self-contained phenotypes i.e. terms, implementation, validation evidence. We only focused on rule-based approaches and did not cover machine learning approaches. While rule-based methods are the most widely used in the UK, data-driven high-throughput approaches including natural language processing methods are emerging [Zhou *et al*., 2016, Pikoula *et al*., 2019]. These approaches pose different challenges and their requirements would need to be documented and analysed in order to ensure their integration [Hripcsak & Albers 2013]. Finally, reproducible research approaches [Denaxas *et al.,* 2017, Goodman *et al,* 2016] which are covered elsewhere would also need to be carefully taken into consideration in order to ensure algorithm portability.

## 6. Steps towards a minimum information standard

Based on our findings, we propose that an EHR phenotyping algorithm representation combines metadata, implementation logic, validation evidence and use-cases.

We suggest the following components towards establishing a minimum information standard with regards to rule-based phenotyping algorithms for UK EHR:

### Part 1 – Algorithm metadata

Succinct information about the goal of the algorithm, the intended use-case, the data sources and controlled clinical terminologies used, applicable age groups and genders, list of authors and their contact details and a set of SNOMED-CT terms to classify the algorithm. A unique identifier, such as a Digital Object Identifier (DOI), should be minted to enable usage tracking in subsequent research.

### Part 2 – Implementation

Details on the implementation logic of the algorithm with pseudocode to facilitate the translation to machine code and documentation on decisions made and reasoning. Where possible analytical scripts should be attached using markdown or a similar approach. The standard should support defining complex Boolean and temporal logic across multiple EHR sources and clinical terminologies. In the future, a computable phenotype format should encapsulate this information as a stand-alone file.

### Part 3 – Validation evidence

Description of the steps taken to support phenotype validity across six categories (aetiological, prognostic, genetic, expert review, cross-source and external population). For each implementation, the number of cases, controls, NPV and PPV values should be reported and the format should support the embedding of graphical files (e.g. forest plots).

### Part 4 – Use-cases

Links to published research utilizing the phenotype algorithms, cross-referenced with DOI’s.

## 7. Conclusion

Our analyses identified a certain level of underlying homogeneity in terms of how phenotyping algorithms are defined and evaluated. We suggest four components towards a minimum information standard that should be used to represent phenotyping algorithms. These findings provide a crucial first step towards curating and disseminating phenotyping algorithms utilizing UK EHR. Further work is required towards establishing a computable format for phenotyping algorithms and ensuring interoperability with other resources (e.g. PheKB).

## Acknowledgments

This work was supported by Health Data Research UK, which receives its funding from HDR UK Ltd (LOND1) funded by the UK Medical Research Council, Engineering and Physical Sciences Research Council, Economic and Social Research Council, Department of Health and Social Care (England), Chief Scientist Office of the Scottish Government Health and Social Care Directorates, Health and Social Care Research and Development Division (Welsh Government), Public Health Agency (Northern Ireland), British Heart Foundation and the Wellcome Trust. The BigData@Heart Consortium is funded by the Innovative Medicines Initiative-2 Joint Undertaking under grant agreement No. 116074. This study was supported by the Farr Institute of Health Informatics Research at UCL Partners (MR/K006584/1). This paper represents independent research part funded by the National Institute for Health Research Biomedical Research Centre at UCLH. HH is a NIHR Senior Investigator. SD is an Alan Turing Fellow.

## Footnotes

1 http://biobank.ndph.ox.ac.uk/showcase/label.cgi?id=42

2 https://www.caliberresearch.org/portal/phenotypes

3 https://www.python.org/

4 http://biobank.ndph.ox.ac.uk/showcase/label.cgi?id=42

5 https://www.caliberresearch.org/portal

6 Primary care EHR available for participants in 2019; case-note review validation underway for multiple phenotypes.

